# The Soil Microbiome of the Caatinga Drylands in Brazil

**DOI:** 10.1101/2024.12.20.629793

**Authors:** Luísa Mayumi Arake de Tacca, Rayane Nunes Lima, Marco Antônio de Oliveira, Patrícia Verdugo Pascoal, Deborah Bambil, Grácia Maria Soares Rosinha, Diana Signor, Marcelo Freire, Elibio Rech

## Abstract

Drylands represent a significant part of the Earth’s surface and include essential and vulnerable ecosystems for the global ecological balance. The Caatinga, with its unique biodiversity adapted to the extreme conditions of this semi-arid region, offers a valuable opportunity to expand our knowledge about these ecosystems. Here, this work reveals the high microbial diversity in the soil and rhizosphere of the Caatinga, with the roots presenting more specialized communities. Bacteria such as *Bacilli*, *Alphaproteobacteria* and *Firmicutes* excelled in critical functions such as nutrient cycling. Interplant differences suggested the influence of root exudates. The metagenomic study of interactions between microorganisms in the rhizosphere of selected plants revealed microbial biodiversity and contributed to our understanding of nutrient cycling, plant growth and resistance to water stress. In addition, they demonstrate biotechnological potential to address global challenges such as desertification and food security.

## INTRODUCTION

Drylands are a widespread and diverse ecological condition present on all continents except Antarctica. They cover more than 47% of Earth’s land surface, with approximately 15% of this area located in Latin America^1,2^. As the largest terrestrial biome, drylands encompass a variety of distinct ecosystems including rangelands, grasslands, woodlands, savannahs, deserts, scrublands, and dry forests^3^. Globally, drylands are defined by several climatic, ecological, and geographical criteria including water deficit, high spatial and temporal variability in precipitation, and seasonal climatic extremes. Precipitation levels categorize drylands into arid, semi-arid, and dry sub-humid regions, with annual rainfall ranging from approximately 10 inches (250 mm) to 30 inches (750 mm)^3,4^. Additionally, drylands are marked by polyextreme conditions such as high evaporation rates, extremely high solar ultraviolet (UV) radiation, and temperature extremes^3,5^. These arid, semi-arid and dry sub-humid environments are marked by extreme climatic conditions, such as irregular precipitation and high evaporation rates, which challenge the adaptation strategies of the plant species and microorganisms that inhabit them. Drylands globally consist of diverse land types, including forests (18%), barren land (28%), grasslands (25%), croplands (14%), and other wooded lands (10%)^6^. Drylands are ecologically significant as they contribute to approximately 40% of global net primary productivity (NPP) and host 35% of the world’s biodiversity hotspots^7,8^.

Furthermore, according to the Food and Agriculture Organization of the United Nations, drylands occupy about 1.1 billion hectares, or 27%, of the global forested area^6^. In South America, drylands predominantly feature arid and semi-arid climates and the primary dryland regions include the Atacama Desert, the Patagonian Desert, the Monte Desert, the Caatinga, and the Chaco region. These areas are critically important for their biodiversity and ecological roles but are also exceptionally vulnerable to climate change and desertification^9^. The Brazilian semi-arid region (SAB) is within this category, with more than twenty-eight million inhabitants, constituting the most populous semi-arid zone in the world^10^. The SAB has a high biodiversity, with the presence of the Caatinga biome (with xerophytic vegetation), along with enclaves of Cerrado (semi-deciduous characteristics), and Atlantic Forest (ombrófila vegetation) regions that create unique ecosystems^11^ and all are characterized by low rainfall rates of <800 mm/year^12–14^. The Caatinga, a unique biome located in the Brazilian semi-arid region, exemplifies the typical characteristics of drylands. With an area of more than 800,000 km², bordered to the east by the Atlantic Ocean and to the west/southwest by the Cerrado and Atlantic Forest Biomes, the Caatinga has a diversity of plants adapted to a climate of low precipitation and high climate variability. However, this biome is still little studied compared to other drylands ecosystems, such as those found in Africa or Asia, which limits the understanding of its ecological dynamics, biodiversity and biological interactions. Additionally, drylands in general host 39% of the global population and as such it is of substantial socioeconomic importance. Recent studies have shown an increase in areas of desertification, plant species becoming more adapted to drought, and an intensification of socioeconomic problems^15,16^.

Drylands support a rich diversity of microorganisms, collectively known as dryland microbiomes, which exist either freely or in symbiotic relationships with vascular plants or within biological soil crusts (biocrusts)^17^. These microbiomes play vital roles in dryland ecosystems, contributing to crucial functions such as the formation of fertile islands, nutrient cycling, and climate regulation, and so are foundational to the ecological succession of vegetation in these extreme environments^3,18^. The Caatinga soil, generally dry and poor in nutrients, is a complex environment of interactions between microorganisms, plants and abiotic factors. The rhizosphere, the region of soil surrounding the plant’s roots, is a dynamic ecosystem where microorganisms play a key role in cycling nutrients, promoting plant growth and mitigating environmental stresses. Symbiotic interactions between the roots of Caatinga plants and microorganisms, especially arbuscular mycorrhizal fungi (AMF), are vital for the adaptation of plants to arid conditions. AMFs, for example, increase the absorption of nutrients and water by plants, essential for their survival in soils with little water availability^19^. Furthermore, AMFs facilitate the acquisition of the essential nutrient phosphorus in poor soils by forming a network of hyphae that expands the area of soil exploration, connecting plant roots to more distant sources of phosphorus, often inaccessible to the roots alone, increasing avaliability^20^. Prokaryotes present in the rhizosphere, such as nitrogen-fixing bacteria, also play a fundamental role in the Caatinga, contributing to nitrogen cycling and improving soil fertility^21,22^. Furthermore, prokaryotes actively participate in the carbon cycle by decomposing organic matter and releasing essential nutrients required for plant growth^21,23^. Another important aspect of the Caatinga soil microbiota is the ability of some bacteria and fungi to mediate the production and consumption of methane, a potent greenhouse gas. The microbial communities present in the soil influence the balance between methane production and oxidation, directly impacting emissions of this gas into the atmosphere^21,24,25^.

Exploring the dryland ecology of microorganisms, its response to climate change and other anthropogenic pressures is nowadays of primary importance. In our study, we collected samples from the soil, rhizosphere and roots of 10 important Caatinga plant species: Umburana (*Commiphora leptophloeos)*, Sapium (*Sapium glandulosum (L.) Morong*), Faveleira (*Cnidoscolus phyllacanthus*), Baraúna (*Schinopsis brasiliensis*), Jurema-preta (*Mimosa tenuiflora)*, Angico (*Anadenanthera colubrina)*, Sete-cascas (*Handroanthus spongiosus)*, Umbuzeiro (*Spondias tuberosa Arruda),* Catingueira *(Caesalpinia pyramidalis Tul.)* and Maniçoba (*Manihot pseudoglaziovii Pax & K. Hoffm.*). These are important plant species in the Brazilian Caatinga and therefore the study of the microbiome composition of such specimens is of high value to the information on the symbioses of microbes in such extreme environments. A map of the Caatinga location in relation to Brazil is shown in Fig.1 as well as a flow diagram with the steps of our study.

**Fig 1:**
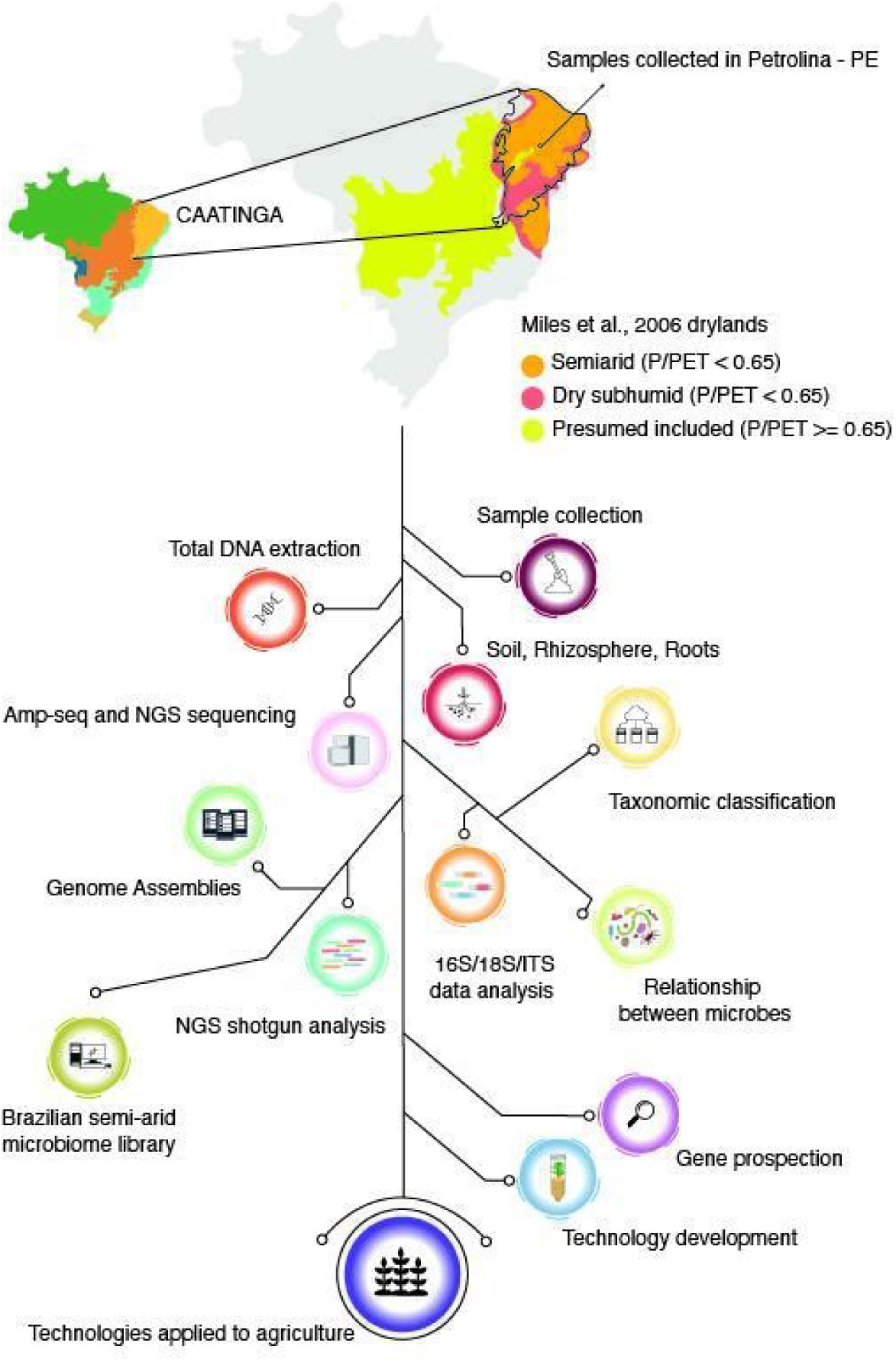
Location of the collected samples and workflow of the SAB study performed.

The harsh conditions of the Caatinga biome, characterized by extreme drought, high temperatures, and soil degradation, create a unique evolutionary pressure that has driven the adaptation of resilient plant species. Therefore, the biotechnological potential of the metagenomic biodiversity of Caatinga microorganisms is considerable. These species have developed mechanisms that help them thrive in challenging environments, and these mechanisms can be harnessed for agricultural benefits, especially through the development of biological products like biopesticides, biofertilizers, and biostimulants. Such biological agents are highly sought due to their low impact on the environment^26,27^. Although the Caatinga biome has been studied in several aspects, the interaction between soil microbiota, plant roots and climatic conditions is still a little explored field. Analyzing their functions and capabilities can reveal new strategies for the development of biotechnological products which can be used in sustainable agriculture, both in the Brazilian semi-arid region and in other regions of the world with dryland characteristics. The use of native microorganisms represents a promising approach to promote more efficient plant growth in arid soils and to protect against disease in agricultural biotechnology^28–31^. The Caatinga, although rich in biodiversity, faces major challenges due to soil degradation, desertification and unsustainable agricultural practices^32,33^. However, exploring microorganism-plant interactions, especially in regions with preserved or recovering native vegetation, can offer new opportunities to restore soil functionality and improve land productivity in the semi-arid region. Studying the rhizosphere microbiota of species typical of the Caatinga, through advanced metagenomic sequencing techniques, not only contributes to the basic understanding of the ecology of the biome, but also opens ways for the use of its biotechnological potential in promoting more sustainable and efficient agricultural practices. Knowledge about the diversity and functions of the microorganisms present in this ecosystem can provide crucial insights for the sustainable management of the Caatinga and its adaptation to climate change. Furthermore, by expanding the study to a metagenomic perspective, it is possible to map the networks of interactions that support the health of the soil and vegetation present, providing valuable tools for the conservation and rational use of this biome of great ecological and economic importance for Brazil and to the world.

## RESULTS

### Rhizosphere and soil share higher diversity and common microbes

We collected 10 plant samples from the Caatinga and sorted these into the soil, rhizosphere, and root regions and the total DNA was isolated according to ^27^. Upon analysis of the 16S regions of the samples, we grouped the results by the previously mentioned categories (here referred to as microbiome sectors). Our goal was to determine how closely related the microbiomes from these three regions of each plant were. We anticipated that soil and rhizosphere samples would share many of the same organisms, while the root samples would exhibit a more distinct microbial community. In Fig. 2a, we can see the differences between these three microbiomes. As expected, the soil and rhizosphere samples show a higher rarefaction curve, indicating greater species richness compared to the roots, which appear lower on the graph. None of the curves reach a plateau, suggesting that deeper sampling would be beneficial in future studies.

**Fig. 2:**
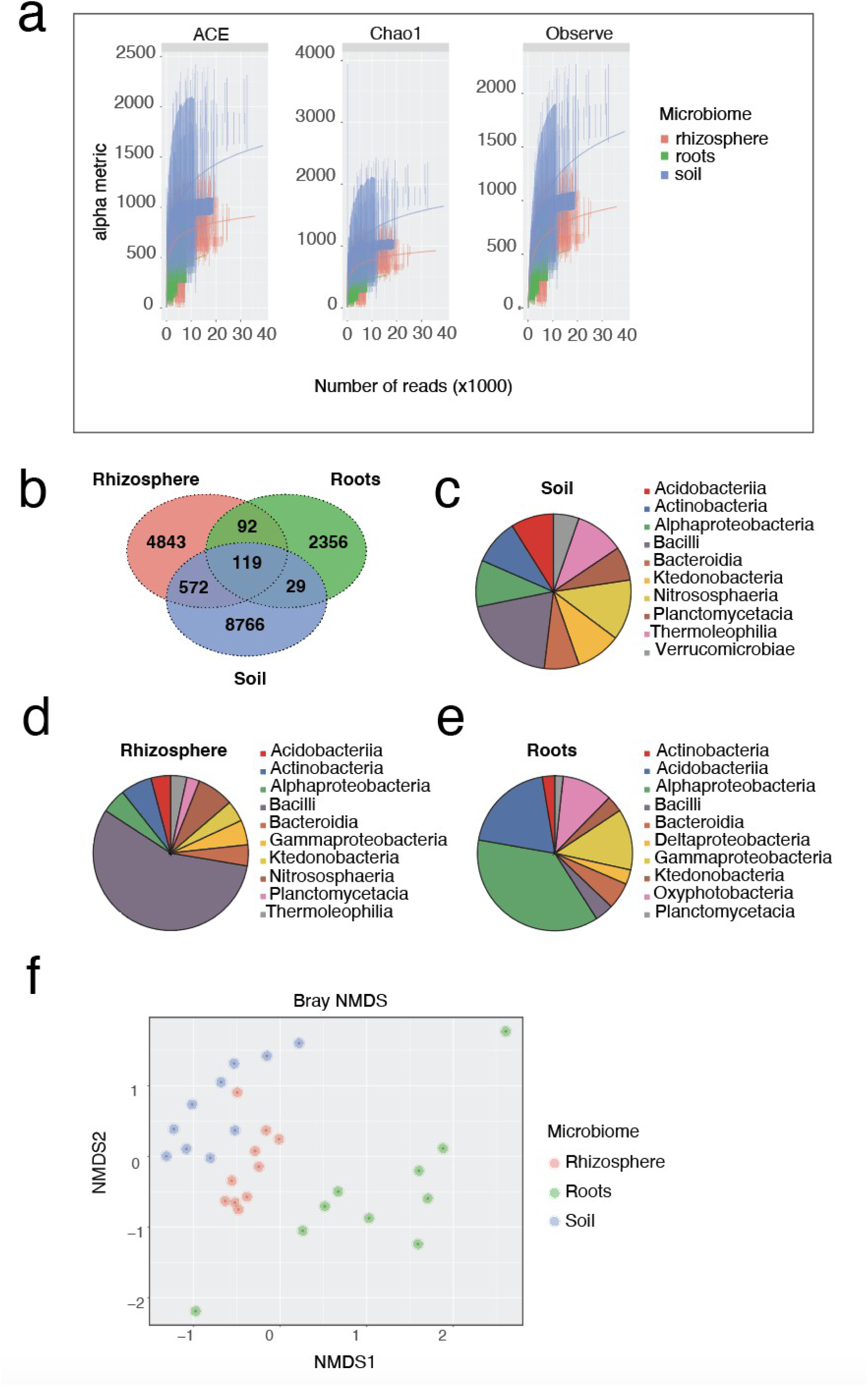
The general microbiome landscape from t h e soil, roots and rhizosphere of the Brazilian semi-arid region Caatinga. **a** Rarefaction curves of the Caatinga samples are separated into roots, soil and rhizosphere. **b** Venn diagram of the OTUs between the roots, soil and rhizosphere of the Caatinga samples. **c, d, e** Pie chart with the top 10 taxa (class) for the soil, rhizosphere and roots of the Caatinga biome respectively. **f** Bray NMDS ordination of the Caatinga samples separated by soil, roots and rhizosphere.

We then used the operational taxonomic units (OTUs) to examine the overlap between the rhizosphere, roots, and soil, which is shown in the Venn diagram (Fig. 2b). Consistent with our expectations, more OTUs were shared between the soil and rhizosphere (572) than between the roots and soil (29) or the roots and rhizosphere (92).

When comparing the landscape of the top 10 classes present in each of these groupings (Fig. 2c-e), we observed that the soil microbiome had a more evenly distributed microbial composition, whereas the roots and rhizosphere were dominated by specific groups: Bacilli in the rhizosphere, and *Alphaproteobacteria* and *Acidobacteriia* in the roots. All three sections share the same top five classes: *Acidobacteria*, *Actinobacteria*, *Alphaproteobacteria*, *Bacilli*, and *Bacteroidia*.

In addition to these five classes, the soil and rhizosphere also share *Ktedonobacteria*, *Nitrososphaeria*, and *Planctomycetacia*. The rhizosphere does not include *Verrucomicrobiae* in its top five classes but does have Gammaproteobacteria, which it shares with the roots. The roots, in contrast, also share *Ktedonobacteria* and *Planctomycetacia* with the soil and rhizosphere, but they additionally contain Deltaproteobacteria and *Oxyphotobacteria*, which are absent from the top five classes from the soil and rhizosphere.

Consistent with our previous observations, the Bray NMDS analysis of the data also shows a distinct separation between the roots and rhizosphere, as well as between the roots and soil, with the rhizosphere lying in between the soil and roots (Fig. 2f).

### Angico shows higher diversity and Baraúna shows lower diversity

After analyzing the microbial patterns in the roots, soil, and rhizosphere, we focused on the plants from which they originated. In this analysis the roots, soil, and rhizosphere were pooled together in favor of the plant of origin (Fig. 3a). We observed that Angico (*Anadenanthera colubrina*) had the highest rarefaction curve, indicating greater richness. On the other hand, Baraúna (*Schinopsis brasiliensis*) had the lowest rarefaction curve and also displayed an undersampling pattern. Consistent with Fig. 2f, we observed a clear separation between the roots, soil, and rhizosphere with the exception of Umbuzeiro (*Spondias tuberosa*), where the soil and rhizosphere showed higher similarity (Fig. 3b). The most distinct soil samples were Jurema-Preta and *Sapium*, as they were on the edges of the soil group. The rhizosphere group was more packed, hinting at a higher similarity between samples while the roots of the plants showed a higher dissimilarity across samples with Umburana and Umbuzeiro on the far edges of the graph. Combining the information from the plants sector microbiomes and the species for each plant, we can see on the heat map (Fig. 3c) there is a high abundance of Proteobacteria in the roots while Firmicutes are more present in soil and rhizosphere but Thaumarchaeota is more present in the soil. As far as the plant species go, Umbuzeiro, Jurema-preta, Caatingueira, Sete-cascas, *Sapium* and Baraúna do not have a high presence of Firmicutes on their roots while Faveleira, Angico, Umburana and Maniçoba do. Umbuzeiro is the only plant that does not have a high presence of Proteobacteria in the roots.

**Fig. 3:**
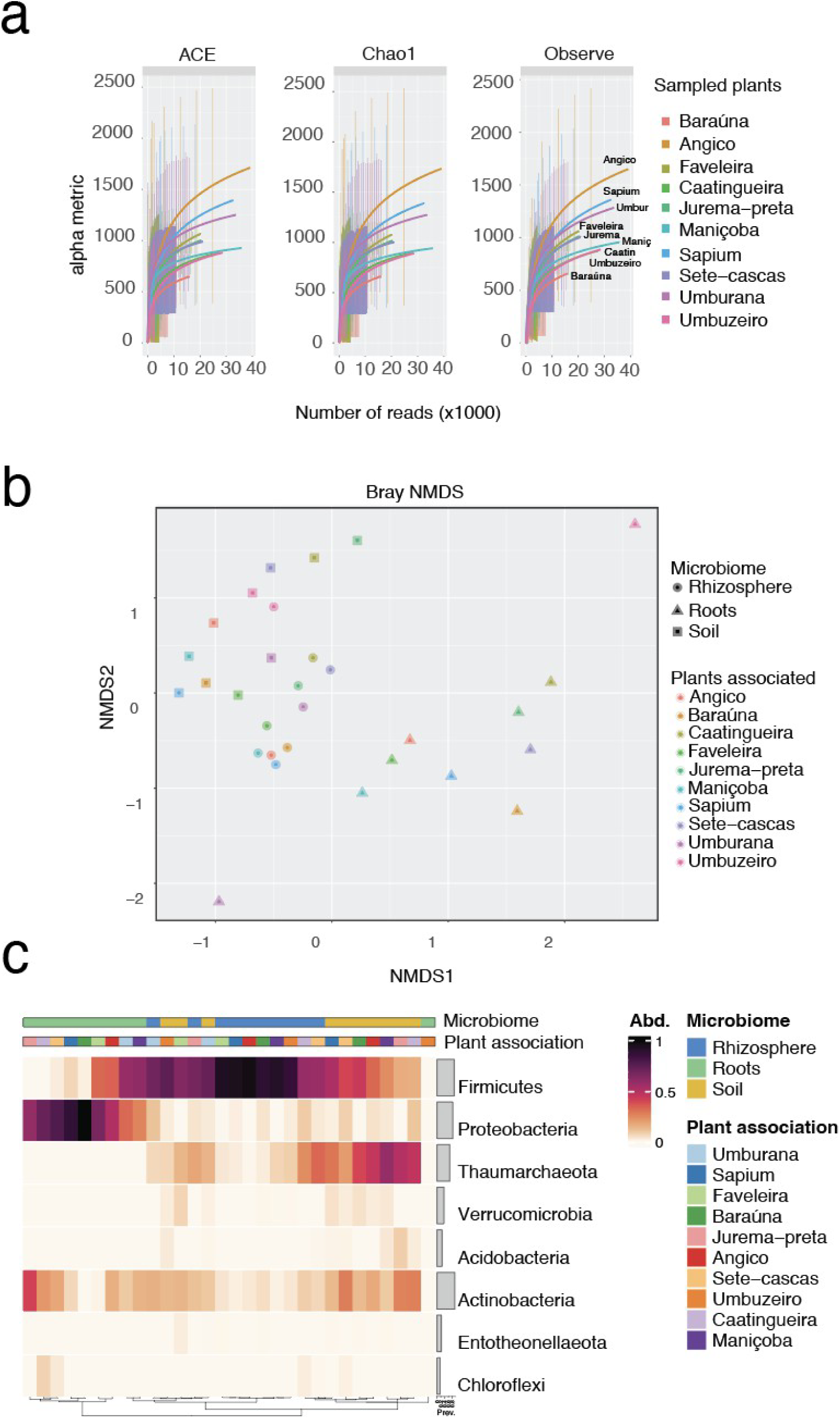
Bacteria landscape associated with the Brazilian sub-arid (Caatinga) typical vegetation. **a** Rarefaction curves of samples associated with 10 different Caatinga plants. **b** Bray NMDS of same samples divided by microbiome (soil, rhizosphere and roots). **c** Heat map of the samples associated with the Caatinga plants and the zoning of the soil, roots and rhizosphere.

### Umburana and Maniçoba show more similarities between each other than with Umbuzeiro

Given the dissimilarities between the roots and the soil and the roots and the rhizosphere, we calculated the Bray-NMDS for the roots separately from the soil and rhizosphere (Fig.4). The pattern of Umburana and Umbuzeiro being the most dissimilar continued in addition to the Maniçoba plant showing a dissimilarity to the rest of the root samples (Fig 4b). Given how the microbes in the roots are important to plant resilience, we sought to understand how these three plants were different from their peers. Checking the overlap between the OTUs, we can observe that Umburana and Maniçoba share more OTUs than Umbuzeiro and Maniçoba or Umburana and Umbuzeiro (Fig. 4c).

**Fig. 4:**
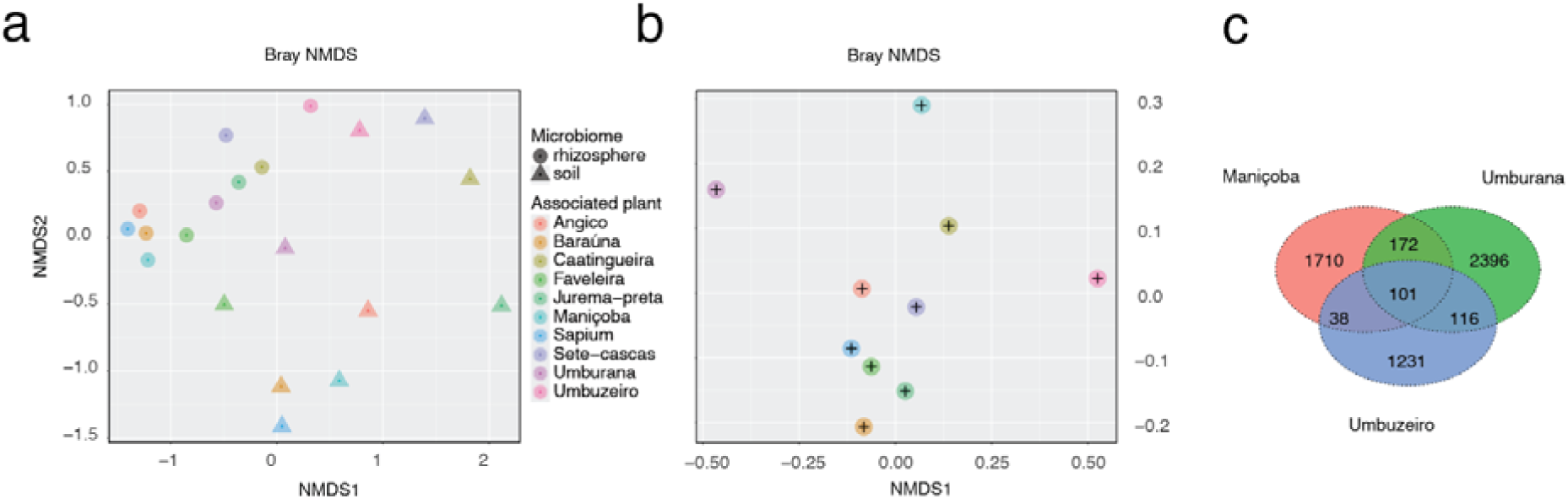
Dissimilarities of the sectors of the Caatinga plants. **a** Bray NMDS of the soil and rhizosphere of the associated microbiome of the Caatinga plants. **b** Bray NMDS of the Caatinga plant soil. **c** Venn diagram of the shared OTUs between Maniçoba, Umburana and Umbuzeiro.

We started by isolating the reads for each plant and by checking the top 8 genera. We then checked if the genera were being exchanged between the soil, roots and rhizosphere for each given plant (Fig. 5). We can see that for Maniçoba (Fig. 5a), in both the soil and rhizosphere, we see the presence of Microseira_Carmichael_Alamaba. There is no overlap between the soil and the roots or between the rhizosphere and roots. On the Umburana plant (Fig. 5b), the overlap is different. Here we see that Kitasatospora is shared between the rhizosphere and the roots, but there is no overlap between the rhizosphere and the soil or between the roots and the soil. On Umbuzeiro (Fig. 5c), there is no overlap between any of the sectors, with all top 8 unique being distinct from each other (Fig. 5 a-c).

**Fig. 5:**
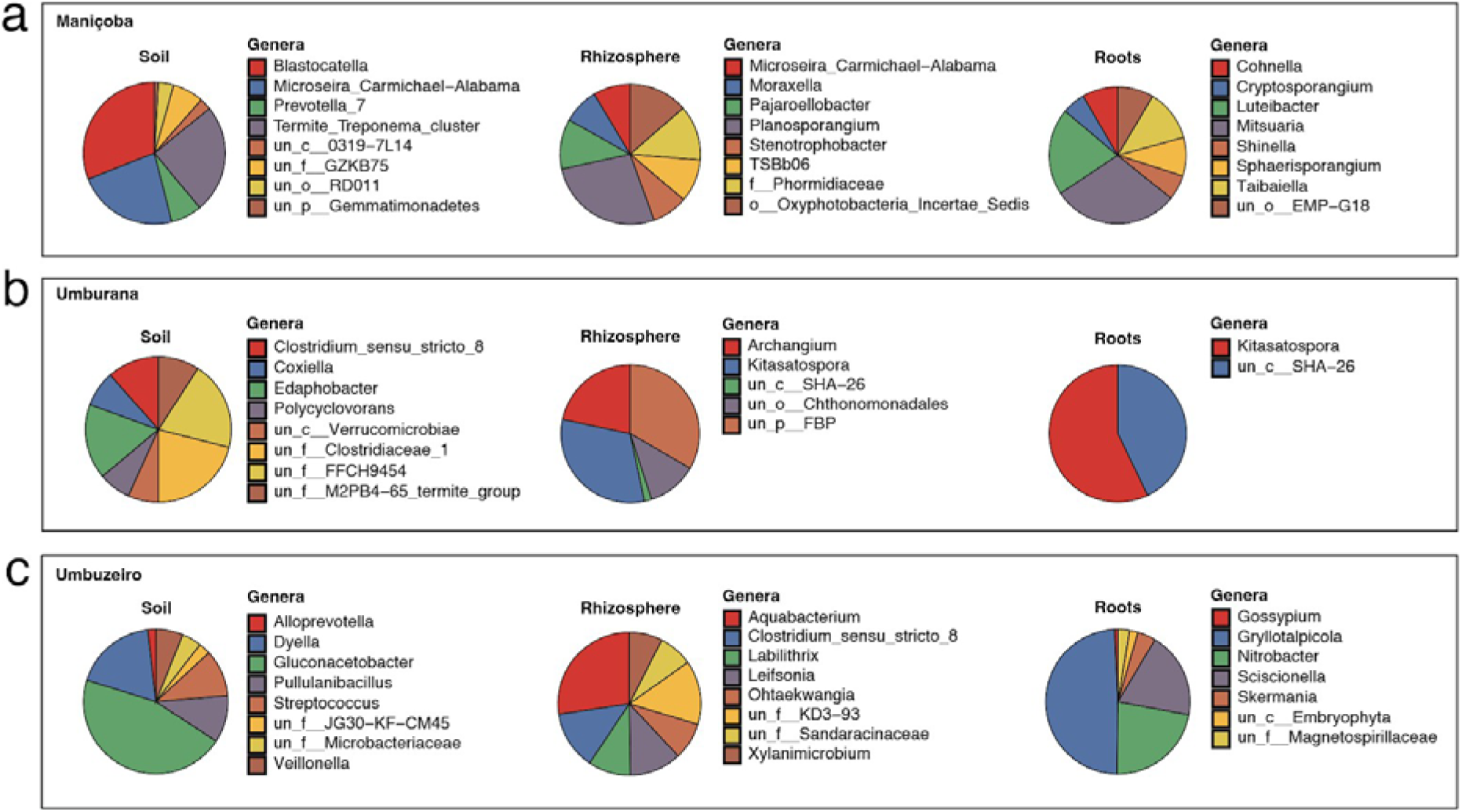
Top 8 unique genera from each plant in the soil, rhizosphere and roots. **a** Maniçoba. **b** Umburana **c** Umbuzeiro

Unique genera were not always the most abundant genera in a given sample. For this reason, we also wanted to determine which were the top 30 most abundant for each of the plant species and each of their sectors. For Umburana, different taxa are prevalent in each sector. The soil had *Bacillaceae* as the most present, followed by *Gemmataceae* and *Chitinophagaceae*. In the rhizosphere, the prevalence was led by *Bacillaceae*, followed by *Burkholderiaceae* and *Solibacteraceae*_subgroup 3. Finally, in the roots, the dominance was not so prominent and *Burkholderiaceae*, *Chitinophagaceae* and an unknown family from order *Acidobacteriales* have similar counts (Fig. 6a).

**Fig. 6:**
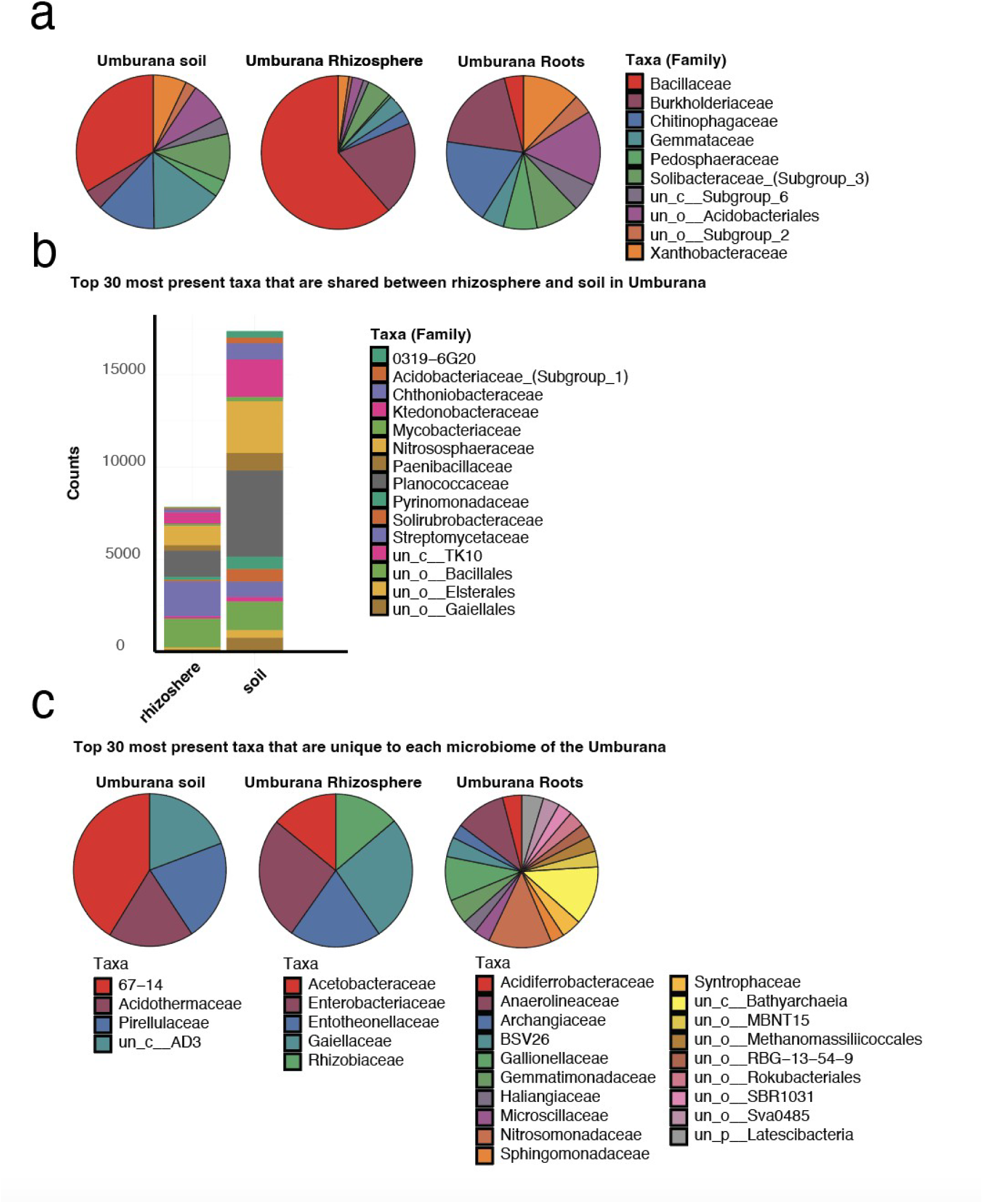
Umburana profile of the top 30 most prevalent taxa shared between. **a** All three sectors and their distribution. **b** two sectors of the Umburana plant. **c** no other sector.

We also checked the taxa that were not common to all sec to rs and only shared between two. We can observe in Fig. 6b that the majority of taxa in this case were shared between the rhizosphere and the soil. From all the shared taxa, the top 5 for the rhizosphere were Streptomycetaceae, an unknown family from *Bacillales*, *Planococcaceae*, *Nitrososphaeraceae* and *Ktedonobacteraceae*. In the soil, *Planococcaceae* was followed by *Nitrososphaeraceae*, *Ktedonobacteraceae*, the unknown family from *Bacillales* and *Paenibacillaceae* (Fig. 6b). Apart from that, there was also an overlap between the roots and the soil regarding the presence of an unknown family from order RCP2-54 that is not shown in the figure. We then checked among the list of the top 30 most abundant taxa, which were unique to either the soil, rhizosphere or roots of that plant. This can be seen in Fig. 6c for Umburana. In the soil, only four of the most abundant taxa were unique to that sector. These were 67-14, *Acidothermaceae*, *Pirellulaceae* and an unknown family from class AD3. The rhizosphere had 5 unique families (*Gaiellaceae*, Enterobacteriaceae, *Entotheonellaceae*, *Acetobacteraceae* and *Rhizobiaceae*) within the top 30 most abundant taxa, they were evenly present. The roots were the sector with the most unique families present in the top 30 taxa. This can be seen on Fig. 6c bottom right.

We did the same study with Umbuzeiro and Maniçoba (Fig. 7 and 8 respectively). The Umbuzeiro had a different pattern of distribution of the common top 30 taxa. There was no dominance of taxa in the soil, however, *Ktedonobacteraceae* was the most present. In the rhizosphere, *Burkholderiaceae* was the most prevalent with 34% of the counts and in the roots, we had a strong *Burkholderiaceae* presence as well, with 80% of the counts (Fig. 7a). Regarding the shared taxa between two sectors, we saw t h a t the rhizosphere and soil shared the most taxa with the most abundant being *Planococcaceae*, an unknown family from order *Bacillales*, *Nitrososphaeraceae* and *Bacillaceae* (Fig. 7b). The rhizosphere shared Caulobacteraceae, *Micropepsaceae*, *Nocardioidaceae* and *Sphingomonadaceae* with the roots but with a lower abundance (Fig. 7c). The roots and the soil had *Acidothermaceae*, an unknown family from order *Betaproteobacteriales* and family 67-14 in common (Fig. 7b). Similarly to what was observed for Umburana, the roots from Umbuzeiro showed the least number of unique, top 30 taxa compared to the soil or rhizosphere. The soil had an evenly spread distribution of *Erysipelotrichaceae*, *Gemmatimonadaceae*, *Pyrinomonadaceae* and an unknown class TK10. The rhizosphere had three unique hits (*Acidobacteriaceae* subgroup 1, *Sphingobacteriaceae* and unknown class subgroup 6) and finally, the roots had 15 unique taxa, with dominance of *Pseudonocardiaceae* (Fig. 7c).

**Fig. 7:**
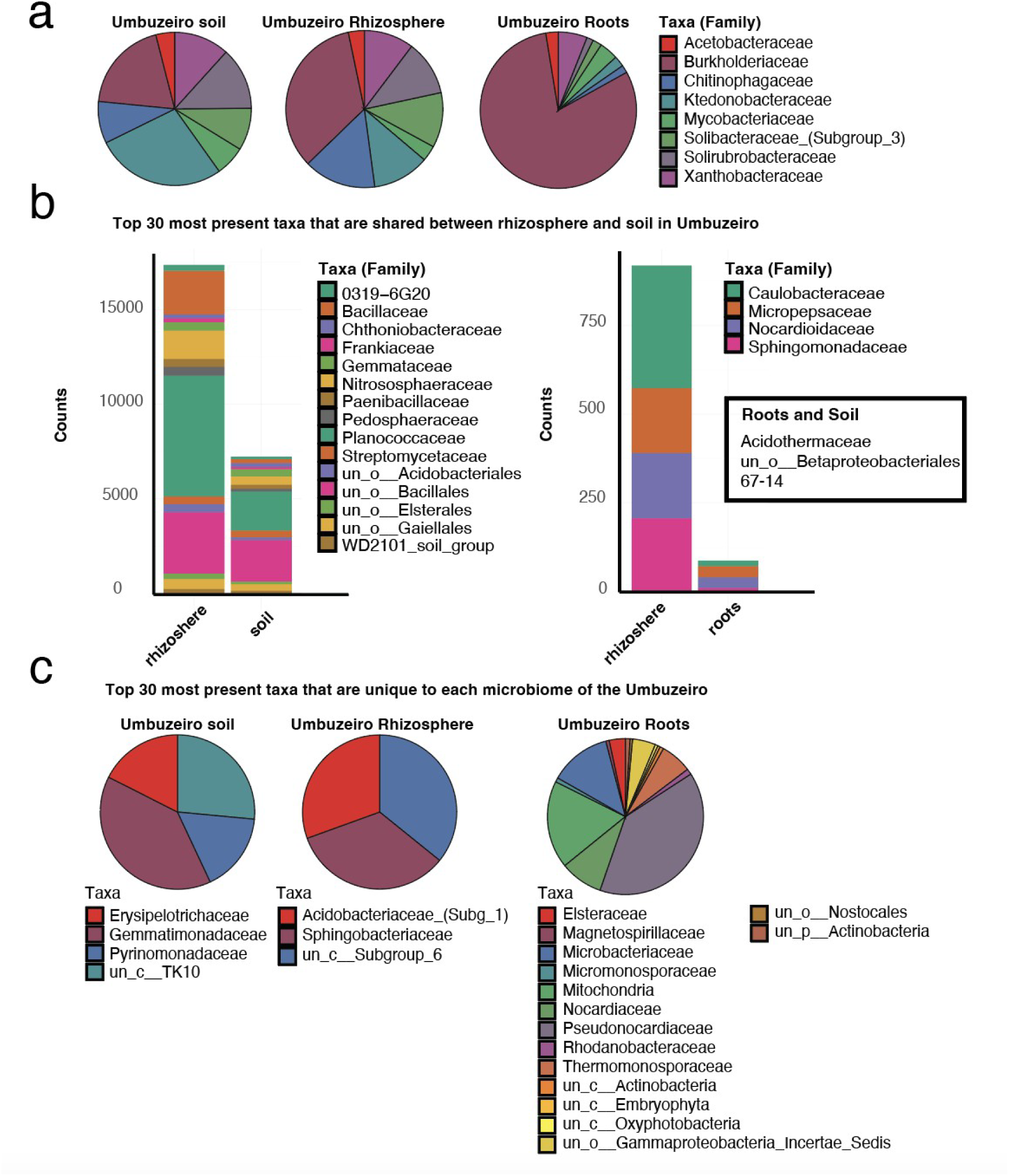
Umbuzeiro profile of the top 30 most prevalent taxa shared between. **a** all three sectors and their distribution. **b** two sectors of the Umbuzeiro plant. **c** no other sector.

**Fig. 8:**
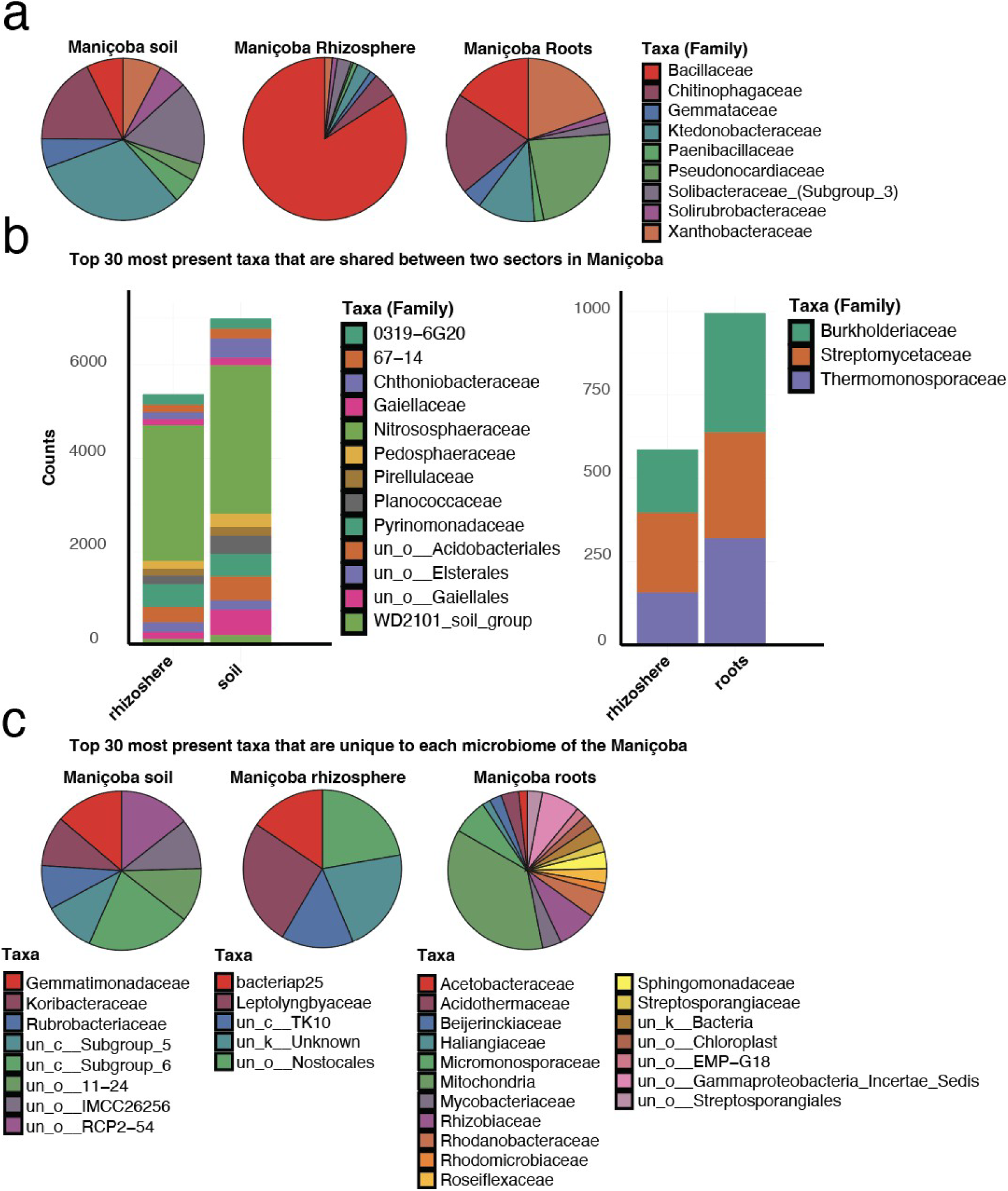
Maniçoba profile of the top 30 most prevalent taxa shared between. **a** all three sectors and their distribution. **b** two sectors of the Maniçoba plant. **c** no other sector.

Maniçoba was the plant with the most distinct pattern (Fig. 8). The top 30 most abundant taxa in the soil was dominated by *Chitinophagaceae*, *Ktedonobacteraceae* and *Solibacteraceae* subgroup 3. The rhizosphere had a clear dominance of *Bacillaceae* while the roots had *Bacillaceae*, *Chitinophagaceae*, *Pseudonocardiaceae* and *Xanthobacteraceae* as the most abundant (Fig. 8a). The rhizosphere and soil were the sectors with the most shared taxa with 13 shared families and a clear dominance of the WD2101 soil group (Fig. 8b). The rhizosphere and roots shared 3 taxa that were evenly distributed (*Burkholderiaceae*, *Streptomycetaceae*, and *Thermomonosporaceae* (Fig. 8b). In comparison to Umbuzeiro and Umburana, Maniçoba had more unique taxa in the top 30; with the soil showing eight unique taxa, rhizosphere showing five unique taxa and roots showing 18 unique taxa (Fig. 8c).

## Discussion

The results confirmed the intimate and distinct relationship between the soil, rhizosphere and root microbiomes in Caatinga, highlighting significant ecological patterns related to microbial diversity and their functions. As expected, the soil and rhizosphere showed greater species richness, evidenced by higher rarefaction curves, while the roots exhibited more specialized microbial communities. This pattern reflects the functional complexity of interactions between plants and microorganisms, which vary depending on the sector, as already reported in studies on arid and semi-arid ecosystems. The greater overlap of OTUs between soil and rhizosphere, when compared to the roots and other sectors, highlighted the functional transition between microbial communities living freely in the soil and those associated with the root surface. Classes such as *Bacilli*, *Alphaproteobacteria* and *Acidobacteriia* dominated the more specialized sectors, suggesting that the rhizosphere and roots favor microbial groups adapted to specific plant-microorganism interaction functions. This specificity was particularly evident in the dominance of *Alphaproteobacteria* in roots, a class known to promote plant growth, cycle nutrients and modulate environmental stresses. Significant differences in patterns of microbial diversity and composition were observed between the plant species analyzed. Angico showed greater species richness in the microbiome, while Baraúna showed a lower diversity, suggesting that intrinsic plant characteristics, such as root exudates and root architecture, play a role in modulating the microbiome. The NMDS dissimilarity analysis reinforced these differences, especially in Umbuzeiro, Umburana and Maniçoba, which exhibited unique patterns of microbial composition in the different sectors, highlighting the ecological variability within the biome. The abundance data of the main microbial families also reinforced the functional complexity of the Caatinga microbiome. For example, the high prevalence of *Burkholderiaceae* in the roots of Umbuzeiro highlighted the potential relevance of this family in the adaptation of plants to the semiarid region, while Firmicutes was more abundant in the soil and rhizosphere, indicating their contribution to general soil stability functions. The predominance of *Thaumarchaeota* in the soil suggested its role in nitrogen cycling, especially in soils with low fertility, such as those in the Caatinga. Analysis of the unique taxa in each sector revealed fundamental differences in the ecological functions performed by these communities. The presence of unique, more diverse taxa in the roots of Umburana, Maniçoba and Umbuzeiro indicated an adaptive specialization that may be linked to the production of specific exudates. These would favor microbial groups with exclusive functions, such as resistance to water stress and growth promotion. On the other hand, the significant overlap between soil and rhizosphere in shared taxa such as *Ktedonobacteraceae* and *Nitrososphaeraceae* pointed to the connective function of the rhizosphere as a link between the soil and root microbiome. The highly distinct pattern of microbial overlap in the three species analyzed in greater detail (Umburana, Umbuzeiro and Maniçoba) was also notable. Umbuzeiro showed little overlap between sectors, suggesting a greater functional segregation between the soil, rhizosphere and root microbiomes. In contrast, Umburana and Maniçoba shared more OTUs between rhizosphere and roots, indicating greater interdependence between these sectors in the microbiota of these plants. These data further reinforced the differential impact of microbial communities on critical ecological processes, such as nutrient cycling and promoting plant resilience. The predominance of *Burkholderiaceae* and *Streptomycetaceae* in the rhizosphere and roots highlighted the central role of these groups in plant adaptation to arid soils. Furthermore, the presence of taxa associated with specific functions, such as nitrogen and carbon metabolism, in the roots and the rhizosphere indicated that the Caatinga microbiota has great biotechnological potential, especially for the development of biofertilizers and sustainable bioproducts. Finally, the absence of plateaus in the rarefaction curves suggests that future studies should adopt deeper sampling strategies, allowing an even more comprehensive characterization of the microbial biodiversity of the Caatinga biome. Additional investigations, focused on functional metagenomics, may reveal new ecological interactions and biotechnological applications, enriching knowledge about dryland ecosystems and promoting sustainable management strategies for the Brazilian semi-arid region.

In drylands, the rhizosphere is recognized as a “hotspot” of microbial diversity, playing essential roles in nutrient cycling and plant resilience to drought stress^34,35^. The greater overlap of OTUs between soil and rhizosphere in the Caatinga reflects a global functional pattern, where the soil provides a diverse microbial reservoir that supports root interactions^2,36^. Groups such as *Alphaproteobacteria*, Firmicutes, and Thaumarchaeota, identified in this study, also have critical functions in other drylands, such as nitrogen fixation, carbon metabolism, and plant growth promotion^37,38^. Furthermore, the interplant differences observed in the Caatinga, such as the microbial specialization in Umburana and Umbuzeiro roots, suggest adaptations related to specific plant exudates, parallel to the plant-microbiota interaction strategies described in global drylands^39,40^. These differences can also be partially explained by the physical proximity between plants. Maniçoba is 30 meters from Umburana and 353 meters from Umbuzeiro, while Umburana and Umbuzeiro are 328 meters away. These distances suggest that the local environment and proximity can influence microbial composition by promoting or limiting the dispersal and sharing of microorganisms between different microbiomes. Maniçoba presented a distinct microbial pattern in the analyzed sectors, with the soil dominated by *Chitinophagaceae*, *Ktedonobacteraceae* and *Solibacteraceae*, groups known for their ability to degrade organic matter and maintain soil stability in drylands^35,36,38^. In the rhizosphere, there was a clear dominance of *Bacillaceae*, microorganisms associated with promoting plant growth and nutrient cycling, especially phosphorus and nitrogen, common characteristics in plant rhizospheres in arid environments^37,41^. In the roots, the largest number of unique taxa was observed (18), with emphasis on *Pseudonocardiaceae* and *Xanthobacteraceae*, microbial groups with important roles in biological nitrogen fixation and the synthesis of bioactive compounds that increase plant tolerance to environmental stress^39,40^.The low overlap between the rhizosphere and roots (3 taxa) reflects the high specificity of the root microbiota, a characteristic also reported for dryland plants in other biomes, where the microbial composition is modulated by exudates and specific symbiotic interactions^36^. Furthermore, Maniçoba’s proximity to Umburana (30 m) and its greater distance from Umbuzeiro (353 m) may influence microbial composition due to limited dispersal and local competition between microorganisms, a pattern observed in semiarid regions such as the Sahel and Australia^35,42^. These results highlight the potential of the Caatinga microbiota not only to deepen ecological knowledge, but also to inspire sustainable management strategies in arid regions around the world^42,43^.

## METHODS

### Collection of samples

A total of 10 samples were meticulously collected, with distribution across the Caatinga. To collect the samples, the upper part of the plant was cut and the soil around the plant was drilled to an approximate depth of 30 cm, cutting and removing the main root, while the secondary and tertiary roots remained with the soil sample. Approximately 200 g of soil was collected and, together with the secondary and tertiary roots, when present, placed in zip lock plastic bags, kept on ice and/or refrigerated until shipping. All collection points were photographed, georeferenced by GPS and described based on the characteristics of the local vegetation (Table 1). The collected samples were meticulously divided into soil, roots and rhizosphere components. The soil samples were manually sieved to remove rocks and roots in order to generate a uniform sample. Then, the rhizosphere was obtained by washing the roots with PBS 1X pH8.3. Finally, the roots were carefully removed, and samples followed by centrifugation at 5000 rpm for 10 minutes.

**Table 1.**
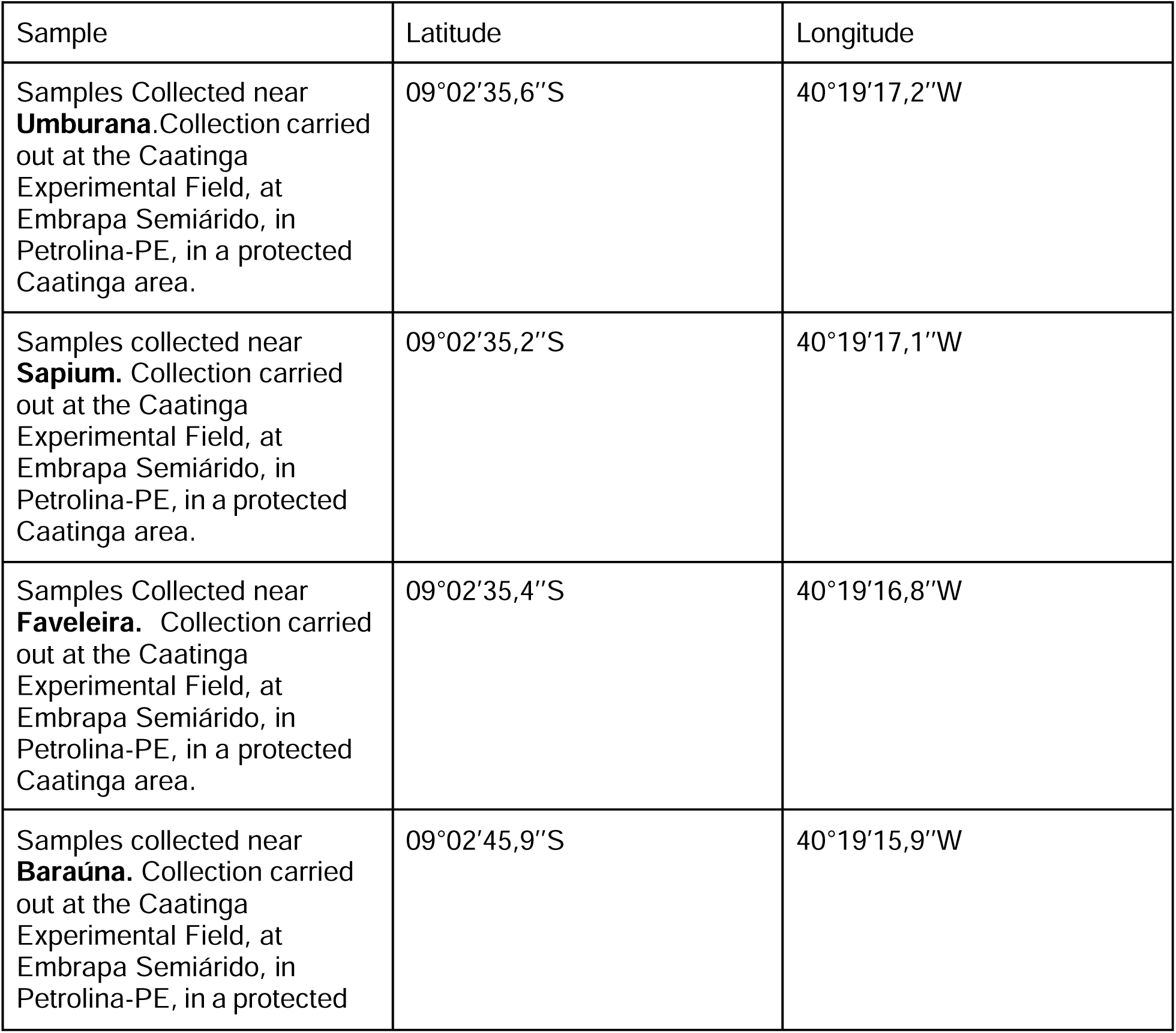
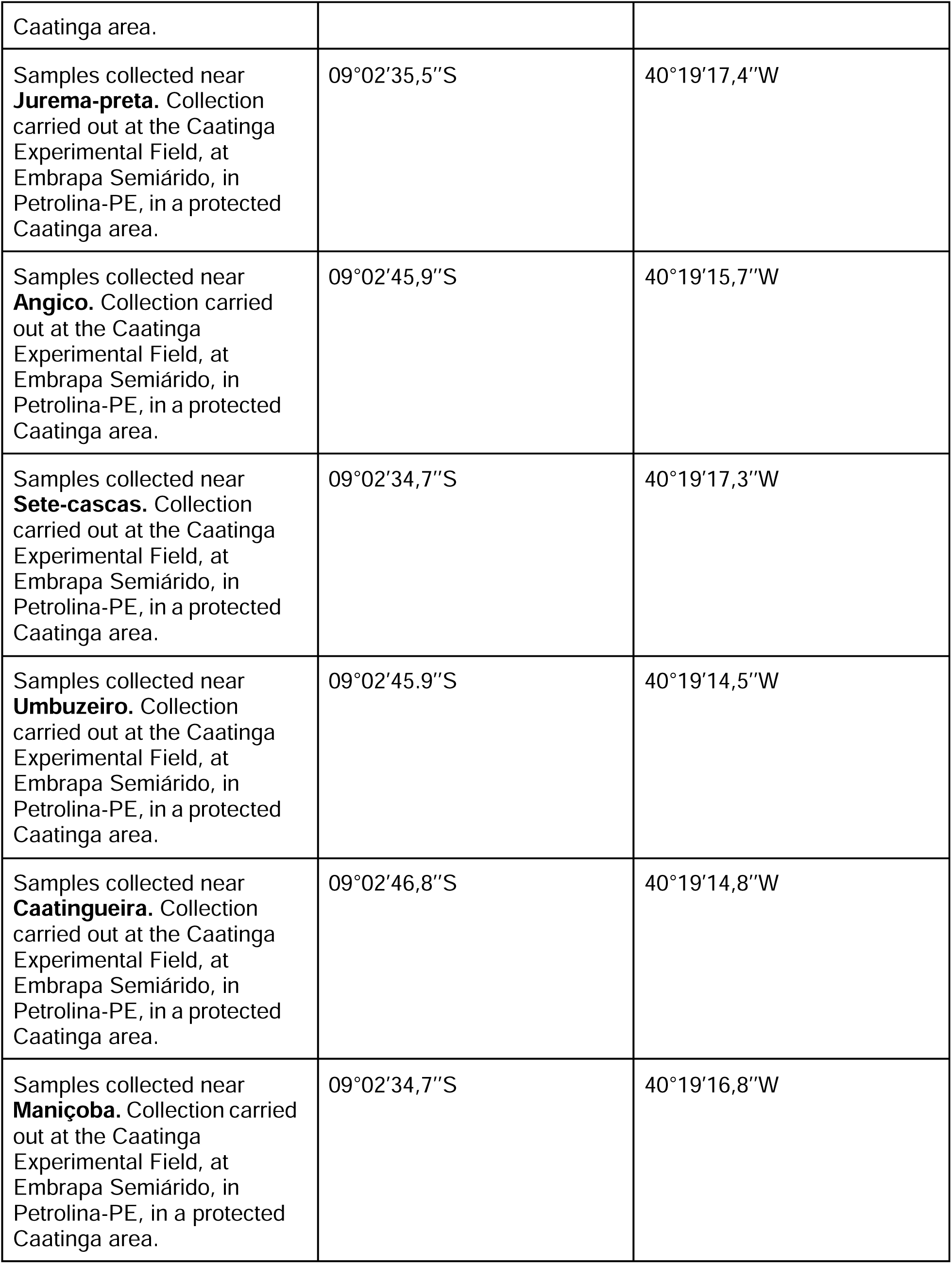
GPS coordinates.

### DNA extraction

The total genomic DNA from the soil and rhizosphere was extracted from 250 mg of each sample using the DNeasy PowerSoil Kit (QIAGEN, Germany) following the manufacturer’s instructions. Plant roots were rinsed in sterile PBS 1x pH 8.3 to remove the rhizosphere and disinfected in a solution of bleach 50% with 0.01% Tween 20 and agitated gently for 3 minutes to remove remaining contaminants. Then, plant roots were then immersed in 70% ethanol for 2 minutes, followed by thorough rinsing with sterile water (bleach/ethanol wash was repeated 3x times). Afterwards, the roots were grounded after freezing with liquid nitrogen followed by total genomic DNA extraction using the The DNeasy Plant Kit (QIAGEN, Germany) following the manufacturer’s instructions. The quality and quantity of DNA were determined by measuring the absorbance at 260/280 nm (A260/A280) on the NanoDrop device (Thermo, Massachusetts, USA) and on Invitrogen™ Qubit™ 4 Fluorometer device (Thermo, Massachusetts, USA), respectively. DNA integrity was verified by 0.8% agarose gel electrophoresis. Sample preparation was done according to McPherson et al. (2018)^44^.

### 16S rRNA gene amplicon library and sequence data analysis

For each area, we used the following 16s primer pair to amplify and sequence this region:

Primers to amplify the 16S V4 region:

515F GTGCCAGCMGCCGCGGTAA and 806R GGACTACHVGGGTWTCTAAT

The amplifications and amplicon-sequencing were performed at JCVI.

### Shotgun sequencing and data processing

For all areas, samples were normalized and pooled. All root, rhizosphere and soil samples for any given area were merged into a single sample. We ran samples on a gel to check for degradation (Supplemental Fig. 1). 50 ng of each pool was sequenced using Illumina shotgun at Novogene.

For the shotgun sequencing data, samples were processed and cleaned by the sequencing company Novogene. Cleaned sequencing files were paired and a library was made for each biome. These libraries were used to classify the microbes using Kaiju through the standard settings in the KBase platform^45^.

For the amplicon-sequencing data, the DADA2 pipeline was utilized to process the clean reads^46^, leading to the formation of operational taxonomic units (OTUs). Taxonomic classification of each representative read and OTU was carried out using the ribosomal database project (RDP) classifier within the SILVA database for bacterial species (with a confidence level of 70%) and the UNITE database for fungal species^47,48^. The analysis of OTUs included their relative abundance at both the genus and phylum levels. Analysis was done using the Phyloseq package^49^.

## FIGURES

Figures were made in Adobe illustrator. References to the package used to calculate the figures in R are described in each figure legend.

## DATA AVAILABILITY

All data generated or analyzed during this study are included in this published article and its supplementary information files. Correspondence and requests for materials should be addressed to Elibio Rech.

## ACKNOWLEDGEMENTS

We thank Magna Soelma Beserra de Moura from Embrapa Semi-arid, Petrolina, Recife, Brazil, for all her assistance with the logistics of sample collection and acquisition.

We also thank the financial support from Embrapa Genetic Resources and Biotechnology/National Institute of Science and Technology in Synthetic Biology, National Council for Scientific and Technological Development/ Ministry of Agriculture Livestock and Supply (465603/2014-9; 400145/2023-5), Research Support Foundation of the Federal District (0193.001.262/2017), and Coordination for the Improvement of Higher Education Personnel.

## AUTHOR CONTRIBUTIONS

Conceptualization: L.M.A.T., R.N.L., and E.R.

Gathered data: G.M.S.R., D.S., and M.F.

Funding acquisition: E.R.

Project administration: E.R.

Supervision: E.R.

Writing—original draft: L.M.A.T., and R.N.L.

Writing – review & editing: L.M.A.T., R.N.L., M.A.O, P.V.P., D.R.B., D.S., M.F. and E.R.

All authors completed, edited and approved the final version.

## References

1 Reynolds, J. F. et al. Global desertification: building a science for dryland development. Science (New York, N.Y.) 316, 847–851 (2007). 10.1126/science.1131634

2 Koutroulis, A. G. Dryland changes under different levels of global warming. The Science of the total environment 655, 482–511 (2019). 10.1016/j.scitotenv.2018.11.215

3 Coleine, C. et al. Dryland microbiomes reveal community adaptations to desertification and climate change. The ISME journal 18 (2024). 10.1093/ismejo/wrae056

4 Selari, P. et al. Short-Term Effect in Soil Microbial Community of Two Strategies of Recovering Degraded Area in Brazilian Savanna: A Pilot Case Study. Frontiers in microbiology 12, 661410 (2021). 10.3389/fmicb.2021.661410

5 Coleine, C., Stajich, J., De los Ríos, A. & Selbmann, L. Beyond the extremes: Rocks as ultimate refuge for fungi in drylands. Mycologia (2020). 10.1080/00275514.2020.1816761

6 FAO. Trees, forests and land use in drylands: the first global assessment. 1 edn, (FAO, 2019).

7 Wang, L. et al. Dryland ecohydrology and climate change: critical issues and technical advances. Hydrology and Earth System Sciences Discussions 9 (2012). 10.5194/hessd-9-4777-2012

8 Davies, J., et al. Conserving Drylands Biodiversity. (2012).

9 Ganem, K. A. et al. Mapping South America’s Drylands through Remote Sensing—A Review of the Methodological Trends and Current Challenges. 14, 736 (2022).

10 (IBGE), I. B. d. G. e. E. Censo Brasileiro de 2010. *IBGE* (2012).

11 de Oliveira Silva, A. K. AB’SÁBER, AZIZ NACIB. OS DOMÍNIOS DE NATUREZA NO BRASIL: POTENCIALIDADES PAISAGÍSTICAS. SÃO PAULO: ATELIÊ EDITORIAL, 2003. Revista de Geografia 29, 252–258 (2012).

12 Silva, M. A. d., et al. Avaliação e espacialização da erosividade da chuva no Vale do Rio Doce, região centro-leste do Estado de Minas Gerais. Revista Brasileira de Ciência do Solo 34 (2010).

13 Marengo, J. A. e. a. in Recursos hídricos em regiões áridas e semiáridas. 383–422 (Campina Grande: Instituto Nacional do Semiárido, 2011).

14 Silva, R. C. et al. The Brazilian semiarid region over the past 21,000 years: Vegetation dynamics in small pulses of higher humidity. Ecological Informatics 77, 102259 (2023). 10.1016/j.ecoinf.2023.102259

15 Silva, R. M. d., Santos, C. A. G., Maranhão, K. U. d. A., Silva, A. M. & Lima, V. R. P. d. Geospatial assessment of eco-environmental changes in desertification area of the Brazilian semi-arid region. Earth Sciences Research Journal 22, 175–186 (2018). 10.15446/esrj.v22n3.69904

16 Oliveira, G. d. C., et al. Soil predictors are crucial for modelling vegetation distribution and its responses to climate change. Science of The Total Environment 780, 146680 (2021). 10.1016/j.scitotenv.2021.146680

17 Song, G., Hui, R., Yang, H., Wang, B. & Li, X. Biocrusts mediate the plant community composition of dryland restoration ecosystems. Science of The Total Environment 844, 157135 (2022). 10.1016/j.scitotenv.2022.157135

18 Wierzchos, J., de los Ríos, A. & Ascaso, C. Microorganisms in desert rocks: the edge of life on Earth. International microbiology : the official journal of the Spanish Society for Microbiology 15, 173–183 (2012). 10.2436/20.1501.01.170

19 Marinho, F. et al. High diversity of arbuscular mycorrhizal fungi in natural and anthropized sites of a Brazilian tropical dry forest (Caatinga). Fungal Ecology 40, 82–91 (2019). 10.1016/j.funeco.2018.11.014

20 Pontes, J. S. et al. Diversity of arbuscular mycorrhizal fungi in Brazil’s Caatinga and experimental agroecosystems. 49, 413–427 (2017). 10.1111/btp.12436

21 Dias, K. C. F. P. et al. Native bacteria from the caatinga biome mitigate the effects of drought on melon (Cucumis melo L.). Comunicata Scientiae 15, e4072 (2023). 10.14295/cs.v15.4072

22 Pinheiro, J. I. et al. Bacterial community in biological soil crusts from a Brazilian semiarid region under desertification process. Scientia Agricola 81 (2024).

23 Costa, D. P. d., et al. Forest-to-pasture conversion modifies the soil bacterial community in Brazilian dry forest Caatinga. Science of The Total Environment 810, 151943 (2022). 10.1016/j.scitotenv.2021.151943

24 Saeed, Q. et al. Rhizosphere Bacteria in Plant Growth Promotion, Biocontrol, and Bioremediation of Contaminated Sites: A Comprehensive Review of Effects and Mechanisms. International journal of molecular sciences 22 (2021). 10.3390/ijms221910529

25 Menezes, R. S. C., Sampaio, E., Giongo, V. & Pérez-Marin, A. M. Biogeochemical cycling in terrestrial ecosystems of the Caatinga Biome. Brazilian Journal of Biology 72 (2012).

26 Araujo, A. S. F. et al. Caatinga Microbiome Initiative: disentangling the soil microbiome across areas under desertification and restoration in the Brazilian drylands. **n/a**, e14298 10.1111/rec.14298

27 Macedo, R. S., Moro, L., Lambais, É. O., Lambais, G. R. & Bakker, A. P. d. EFFECTS OF DEGRADATION ON SOIL ATTRIBUTES UNDER CAATINGA IN THE BRAZILIAN SEMI-ARID. Revista Árvore 47 (2023).

28 Al-Tammar, F. K. & Khalifa, A. Y. Z. Plant growth promoting bacteria drive food security. Brazilian Journal of Biology 82 (2022).

29 Antoszewski, M., Mierek-Adamska, A. & Dąbrowska, G. B. The Importance of Microorganisms for Sustainable Agriculture-A Review. Metabolites 12 (2022). 10.3390/metabo12111100

30 Li, Y. et al. Biofilms formation in plant growth-promoting bacteria for alleviating agro-environmental stress. Science of The Total Environment 907, 167774 (2024). 10.1016/j.scitotenv.2023.167774

31 Chatchatnahalli Tharanath, A., Upendra, R. & Rajendra, K. Soil Symphony: A Comprehensive Overview of Plant–Microbe Interactions in Agricultural Systems. Applied Microbiology 4, 1549–1567 (2024). 10.3390/applmicrobiol4040106

32 Franca Rocha, W. J. S., et al. Towards Uncovering Three Decades of LULC in the Brazilian Drylands: Caatinga Biome Dynamics (1985–2019). 13, 1250 (2024).

33 Lucena, M. S. d., Zakia, M. J. B. & Guerin, N. Discourses on sustainable forest management in the Caatinga Domain. Ambiente & Sociedade 26 (2023).

34 de Queiroz, L. P., Cardoso, D., Fernandes, M. F. & Moro, M. F. in *Caatinga: The Largest Tropical Dry Forest Region in South America* (eds José Maria Cardoso da Silva, Inara R. Leal, & Marcelo Tabarelli) 23–63 (Springer International Publishing, 2017).

35 Huang, J. et al. Global desertification vulnerability to climate change and human activities. 31, 1380–1391 (2020). 10.1002/ldr.3556

36 Rashid, M. I. et al. Bacteria and fungi can contribute to nutrients bioavailability and aggregate formation in degraded soils. Microbiol Res 183, 26–41 (2016). 10.1016/j.micres.2015.11.007

37 Chillo, V., Ojeda, R., Capmourteres, V. & Anand, M. Functional diversity loss with increasing livestock grazing intensity in drylands: The mechanisms and their consequences depend on the taxa. Journal of Applied Ecology 54 (2016). 10.1111/1365-2664.12775

38 Wang, Z. et al. Coupling of Phosphorus Processes With Carbon and Nitrogen Cycles in the Dynamic Land Ecosystem Model: Model Structure, Parameterization, and Evaluation in Tropical Forests. Journal of Advances in Modeling Earth Systems 12, e2020MS002123 (2020). 10.1029/2020MS002123

39 Toby Pennington, R., Prado, D. E. & Pendry, C. A. Neotropical seasonally dry forests and Quaternary vegetation changes. Journal of Biogeography 27, 261–273 (2000). 10.1046/j.1365-2699.2000.00397.x

40 Rito, K., Arroyo-Rodríguez, V., Queiroz, R., Leal, I. & Tabarelli, M. Precipitation mediates the effect of human disturbance on the Brazilian Caatinga vegetation. Journal of Ecology (2017). 10.1111/1365-2745.12712

41 Wu, L., Weston, L. A., Zhu, S. & Zhou, X. Editorial: Rhizosphere interactions: root exudates and the rhizosphere microbiome. 14 (2023). 10.3389/fpls.2023.1281010

42 Martínez-Valderrama, J., Guirado, E. & Maestre, F. T. Desertifying deserts. Nature Sustainability 3, 572–575 (2020). 10.1038/s41893-020-0561-2

43 Durán, P. et al. Microbial Interkingdom Interactions in Roots Promote Arabidopsis Survival. Cell 175, 973–983.e914 (2018). 10.1016/j.cell.2018.10.020

44 McPherson, M. R., Wang, P., Marsh, E. L., Mitchell, R. B. & Schachtman, D. P. Isolation and Analysis of Microbial Communities in Soil, Rhizosphere, and Roots in Perennial Grass Experiments. Journal of visualized experiments : JoVE (2018). 10.3791/57932

45 Arkin, A. P. et al. KBase: The United States Department of Energy Systems Biology Knowledgebase. Nature biotechnology 36, 566–569 (2018). 10.1038/nbt.4163

46 Callahan, B. J. et al. DADA2: High-resolution sample inference from Illumina amplicon data. Nature methods 13, 581–583 (2016). 10.1038/nmeth.3869

47 Ikeda, S., Shimizu, A., Shimizu, M., Takahashi, H. & Takenaka, S. Biocontrol of black scurf on potato by seed tuber treatment with Pythium oligandrum. Biological Control 60, 297–304 (2012). 10.1016/j.biocontrol.2011.10.016

48 Kõljalg, U. et al. UNITE: a database providing web-based methods for the molecular identification of ectomycorrhizal fungi. The New phytologist 166, 1063–1068 (2005). 10.1111/j.1469-8137.2005.01376.x

49 McMurdie, P. J. & Holmes, S. phyloseq: an R package for reproducible interactive analysis and graphics of microbiome census data. PloS one 8, e61217 (2013). 10.1371/journal.pone.0061217

